# A fluctuation-based approach to infer kinetics and topology of cell-state switching

**DOI:** 10.1101/2022.03.30.486492

**Authors:** Michael Saint-Antoine, Ramon Grima, Abhyudai Singh

## Abstract

In the noisy cellular environment, RNAs and proteins are subject to considerable stochastic fluctuations in copy numbers over time. As a consequence, single cells within the same isoclonal population can differ in their expression profile and reside in different phenotypic states. The dynamic nature of this intercellular variation, where individual cells can transition between different states over time makes it a particularly hard phenomenon to characterize. Here we propose a novel fluctuation-test approach to infer the kinetics of transitions between cell states. More specifically, single cells are randomly drawn from the population and grown into cell colonies. After growth for a fixed number of generations, the number of cells residing in different states is assayed for each colony. In a simple system with reversible switching between two cell states, our analysis shows that the extent of colony-to-colony fluctuations in the fraction of cells in a given state is monotonically related to the switching kinetics. Several closed-form formulas for inferring the switching rates from experimentally quantified fluctuations are presented. We further extend this approach to multiple cell states where harnessing fluctuation signatures can reveal both the topology and the rates of cell-state switching. In summary, our analysis provides a powerful approach for dissecting cell-state transitions based on a *single* time point measurement. This is especially important for scenarios where a measurement involves killing the cell (for example, performing single-cell RNA-seq or assaying whether a microbial/cancer cell is in a drug-sensitive or drug-tolerant state), and hence the state of the same cell cannot be measured at different time points.

## I. Introduction

Advances in single-cell technologies have exposed remarkable differences in phenotype and expression patterns between individual cells within the same isogenic cell population [1]–[9]. While some of this variation can be linked to extrinsic factors (i.e., cell-cycle stage, cell size, local extra-cellular environment), increasing evidence points to the role of stochastic processes inherent to gene expression in driving random fluctuations (noise) in gene product levels [10]– [24]. This intercellular phenotypic heterogeneity can play important functional roles in diverse biological processes, from driving genetically-identical cells to different cell fates [25]–[35] to allowing microbes and cancer cells to hedge their bets against uncertain environmental changes [36]–[47]. While single-cell sequencing tools can probe phenotypic heterogeneity within a given cell population, they only provide a static picture of different cell states. Characterizing the dynamics of individual cells transitioning between different states with multi-generational time scales remains a fundamental challenge in advancing the field of single-cell biology. The Luria-Delbrück experiment, also called the “Fluctuation Test”, demonstrated that genetic mutations arise randomly in the absence of selection – rather than in response to selection – and led to a Nobel Prize [48]. We leverage a similar fluctuation-style assay to infer switching dynamics between cellular states. The proposed methodology relies on first growing single cells into colonies and then measuring each colony’s cell-state composition (i.e, the number of cells in different cell states). As a simple example, drug treatment of a single-cell derived colony of cancer cells can classify individual cells into two phenotypic states: drug-sensitive (drug-tolerant) cells that die (survive) in response to treatment. Repeating this process for several colonies yields a distribution for the number of surviving cells. How can we exploit this measured distribution to understand the interchange between cell states?

Starting with a system of reversible switching between two cell states, we derive three different analytical formulas employing different approximations to quantify fluctuations in cell-state composition across colonies and benchmark them with exact stochastic simulations of the cell proliferation/switching process. From a practical standpoint, these formulas are a valuable tool to back-calculate the switching rates from experimentally measured fluctuations at a *single* time point of colony expansion. In the context of the cancer example, inferring the timescale of switching between drug-sensitive and drug-tolerant states is critical for the design of optimal drug treatment schedules [49]. Later on, we further generalize these results to multiple cell states with arbitrary switching topologies. We start by reviewing the original Luria-Delbrück experiment done 80 years ago.

## II. Revisiting the classical Luria-DelbrÜck experiment

By the early 20th century it was known that bacteria can acquire resistance to infection by phages (bacterial viruses). However, it was debated whether mutations leading to resistance were directly induced by the virus (Lamarckian theory), or if they arose randomly in the population before viral infection (Darwinian theory). To discriminate between these alternative hypotheses, Luria & Delbrück designed an elegant experiment where single cells were isolated and grown into colonies. After allowing the colonies to grow for some duration, they were infected by the T1 phage, and the number of resistant bacteria were counted across colonies. If mutations are virus-induced (i.e., no genetic heritable component to resistance), then each bacterium has a small and independent probability of acquiring resistance, and the colony-to-colony fluctuations in the number of resistant cells should follow Poisson statistics. In contrast, if mutations occur randomly before viral exposure, then the number of surviving bacteria will vary considerably across colonies depending on when the mutation arose in the colony expansion (Fig. 1). The data clearly showed a non-Poissonian skewed distribution for the number of resistant bacteria, validating the Darwinian theory of evolution [48].

**Fig. 1.**
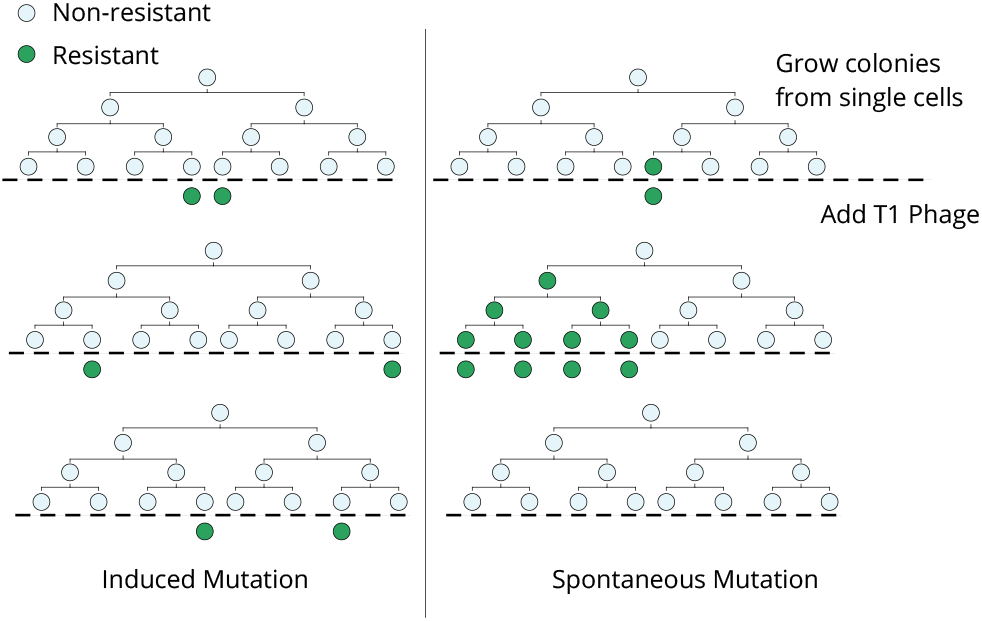
The Luria-Delbrück fluctuation test. Single cells are expanded into colonies and then infected by bacteriophage T1. If resistance mutations are virus-induced, then the number of resistant cells would follow a Poisson distribution across colonies. In contrast, if mutant cells arise spontaneously prior to viral exposure, then there will be considerable colony-to-colony fluctuations in the number of surviving cells, including “jackpot” populations where mutations happened early in the lineage expansion causing many cells to be resistant.

The Luria-Delbrück experiment that came to be known as the “Fluctuation Test”, not only addressed a fundamental evolutionary question leading to a Nobel Prize, but also laid the foundations for the field of bacterial genetics. Apart from its biological significance, the fluctuation test exemplifies the usage of stochastic analysis for uncovering hidden processes even though the underlying cell states may not be directly observable. Publication of the Luria-Delbrück article in 1943 catalyzed rich theoretical work deriving probability distributions for the number of resistant cells based on different biological assumptions [50]–[52], and led to statistical methods for estimating mutation rates from fluctuations in the data [53]–[55]. We refer the reader to [56] for an excellent review of mathematical developments related to the Luria-Delbrück experiment.

The fluctuation test was recently used to study cancer drug resistance [57]. Individual melanoma cells were isolated from a clonal cell population by single-cell FACS sorting, and then grown into colonies. After allowing single cells to expand for a few weeks, the colonies were treated with a targeted cancer drug, vemurafenib. Intriguingly, the colony-to-colony fluctuations in the number of surviving cells were significantly larger than a Poisson distribution, but an order of magnitude smaller than what is predicted by the mutation model. Subsequent analysis showed that stochastic expression of several resistant markers drives individual cells to reversibly switch between drug-sensitive and drug-tolerant states prior to drug exposure [58], and the latter state can transform into a stably-resistant state in the presence of the drug [57]. We next present the mathematical framework for modeling cell-state transitions in an expanding cell colony, followed by the derivation of formulas quantifying the extent of fluctuations as a function of the switching rates.

## III. Fluctuation-approach to infer cell-state switching

Consider a scenario as in Fig. 2 where cells within a population can reside in two states (State 1 & 2). Cells proliferate and reversibly switch between states, and the rates of switching determine the transient heritability of a state, i.e., how many generations it takes to switch back to the other state. Let 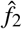 denote the average fraction of cells in State 2 in the original population. Single cells are randomly drawn from the population (through serial dilutions or FACS sorting), and expanded into colonies. Note that the state of the starting single cell is unobservable, as we only consider a *single* endpoint measurement. After growing the colonies for a certain duration of time, each colony is assayed for the fraction of cells in State 2 (or State 1). The basic idea is that if switching between states is relatively fast (several switches happen in the growth duration), then the fraction of State 2 cells will rapidly equilibrate to 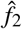 in each colony, and colony-to-colony fluctuations will be minimal (Fig. 2). In contrast, if switching is slow, then based on the memory of the initial cell, colonies will primarily be composed of cells in either State 1 or 2 generating large colony-to-colony fluctuations (Fig. 2). **In essence, fluctuations in colony cell-state composition reveals the timescale of switching, with slower relaxation kinetics driving higher fluctuations**.

**Fig. 2.**
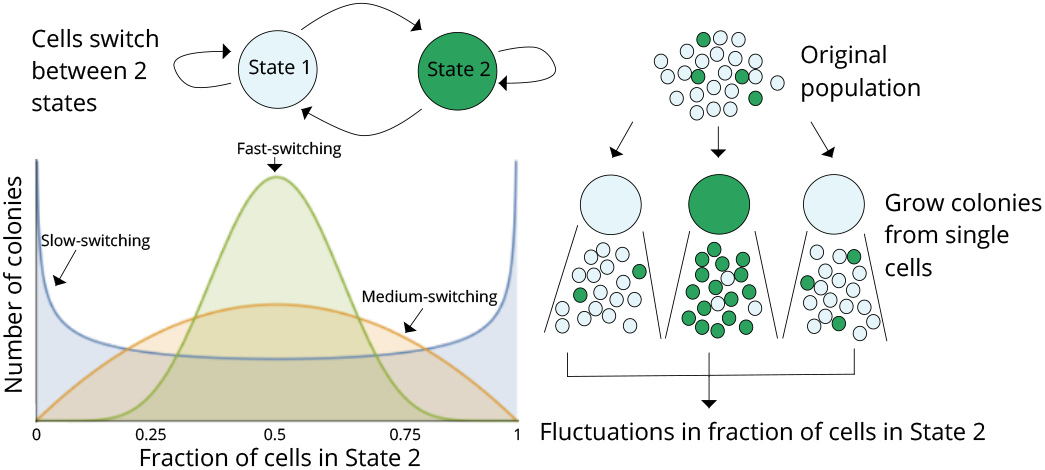
A fluctuation-test approach for deciphering switching between two cellular states. Schematic showing cells in two different states (State 1 & 2) together with reversible switching between states, and proliferation in each state. Individual cells are randomly chosen from the original population and assayed for the fraction of cells in State 2 after a certain duration of lineage expansion. If switching between states is relatively fast, then colonies will show similar fractions of State 2 cells as the original population, and variance across colonies will be minimal. On the other hand, if switching is slow, then colony composition will heavily depend on the state of the initial cell, and there would be large colony-to-colony fluctuations based on differences in the initial condition. Thus, statistical fluctuations in colony composition can be exploited to infer the transient heritability of cellular states.

To directly relate these fluctuations to the switching kinetics, we model the cell-colony expansion process by considering that cells proliferate with a constant rate *k*_*x*_ implying a mean cell-cycle time of 1*/k*_*x*_ (i.e., time taken by each cell to finish cell-cycle and make two daughters). Cells in State 1 transition to State 2 with a constant rate *k*_1_, and switch back to State 1 with a constant rate *k*_2_ resulting in the average fraction

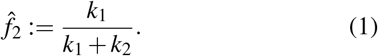

We make several simplifying assumptions in our model formulation:

- The proliferation rate of a cell is the same irrespective of the cellular state.
- There is no cell death in the expanding colony.
- The population remains in the exponential growth phase during the time span of the experiment.
- The switching rates are independent of the number of cells in the colony.

Assuming that the starting cell at time *t* = 0 is chosen randomly from the original population, its state follows a Bernoulli random variable: the cell is in State 2 with probability 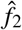 and in State 1 with probability 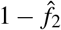. Let stochastic processes *x*_1_(*t*) and *x*_2_(*t*) denote the number of cells in State 1 and 2, respectively, with total cell number *x*(*t*) = *x*_1_(*t*) + *x*_2_(*t*). Our goal is to quantify colony-to-colony fluctuations in the fraction of cells in State 2

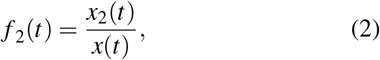

at time *t* of colony expansion. It is important to point out that we work with the fraction of cells in State 2, and not the number of cells in State 2. The rationale for this is that fluctuations in the former appear more robust to the cell-cycle time distribution. For example, stochastic simulations of an expanding colony with an exponentially-distributed or a lognormally-distributed cell-cycle time (with the same mean) result in different levels of fluctuations in *x*_2_(*t*), but similar fluctuations in *f*_2_(*t*) (Fig. 3). Employing different assumptions, we next derive closed-form formulas for the coefficient of variation (standard deviation over mean) of *f*_2_(*t*) across colonies.

**Fig. 3.**
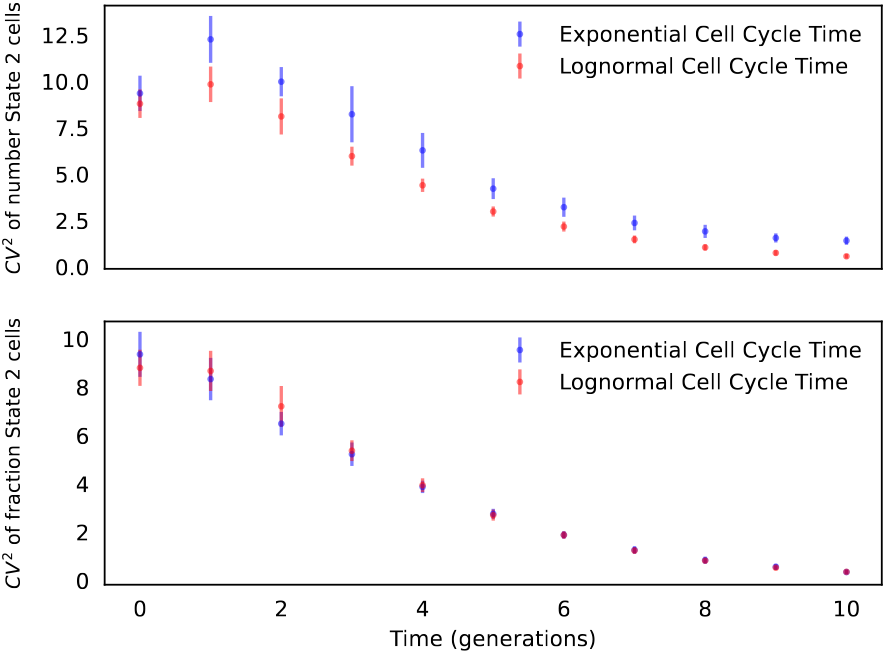
Fluctuations in state fractions are robust to the cell-cycle time distribution. Fluctuation in the number (top) and the fraction (bottom) of cells in State 2 as a function of colony-expansion time. Here and in all other figures we set the mean cell-cycle time to 1 time units by having *k*_*x*_ = 1, and *t* can then be interpreted as time in the number of cell generations. Colony growth simulation was done assuming two different cell-cycle distributions: exponentially-distributed and lognormally-distributed cell-cycle time with mean 1 and variance 0.28. Other parameters were taken as 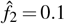, *k*_2_ = 1*/*5 and the coefficient of variation was computed across 1000 colonies (i.e., simulation runs). These simulations were then repeated 20 times to generate error bars that show one standard deviation.

### A. Deterministc proliferation approximation

Ignoring any stochasticity in the cell proliferation and switching processes, colony growth from a single cell can be modeled as an ordinary differential equation

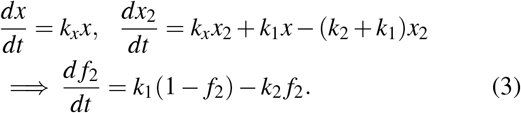

Solving (3) for a Bernoulli initial condition *f*_2_(0) = 1 with probability 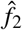(starting cell is in State 2) and *f*_2_(0) = 0 with probability 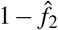 (starting cell is in State 1) yields

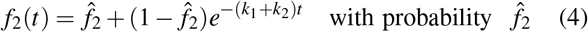

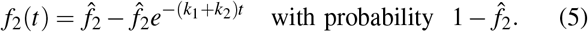

This results in the following mean fraction

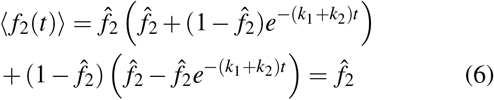

where ⟨ ⟩denotes the expected value of a random process. As intuitively expected, at any time in the lineage expansion the mean fraction of cells in State 2 across colonies is the same as that in the original population. Using a similar approach for the second-order moment

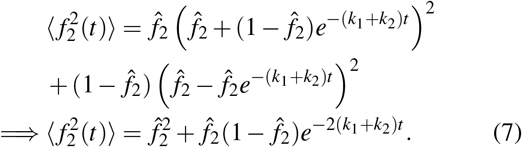

This leads to the following colony-to-colony fluctuations

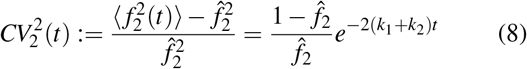

with *CV*_2_ being the coefficient of variation (*CV*) of the State 2 fraction. The formula shows that for a fixed 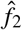 and time *t*, faster switching attenuates fluctuations with *CV*_2_ → 0 as *k*_1_, *k*_2_ → ∞. By defining a dimensionless quantity

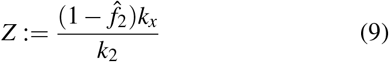

we can rewrite (8) as

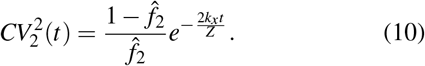

Given a time *t* of *CV*_2_ measurement, switching rates can be inferred by simultaneously solving (1) and (10) based on a-priori knowledge of the cell proliferation rate, and 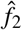 is the average fraction of State 2 cells across colonies.

### B. Stochastic proliferation model

To capture low-copy number effects at early stages of colony expansion, we consider a stochastic model formulation where now *x*(*t*), *x*_1_(*t*), *x*_2_(*t*) *∈* {0, 1, 2, …} are integer-valued random processes. For defining the model and its subsequent analysis we work with *x*(*t*) and *x*_2_(*t*). The time evolution of these populations is governed by the events in Table 1 that occur probabilistically with rates given in the third column. Whenever the event occurs, the corresponding reset map is activated and cell numbers increase/decrease by one. This formulation corresponds to the cell-cycle time being drawn from an exponential distribution with mean 1*/k*_*x*_. Moreover, the time spent in States 1 and 2 is exponentially-distributed with means 1*/k*_1_ and 1*/k*_2_, respectively.

**TABLE I.**
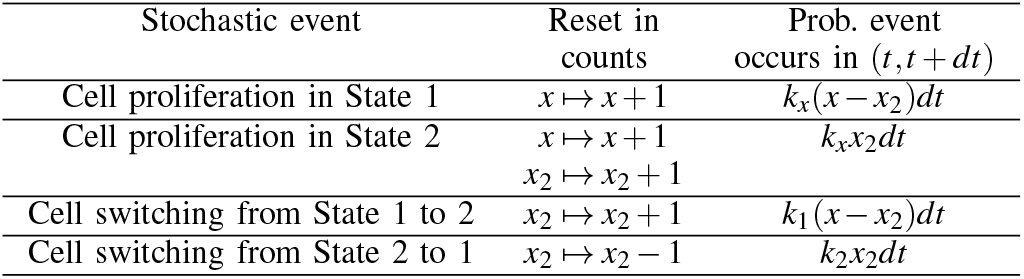
Stochastic model of cell proliferation and switching.

| Stochastic event | Reset in counts | Prob. event occurs in $(t, t+dt)$ |
| --- | --- | --- |
| Cell proliferation in State 1 | $x \mapsto x+1$ | $k_x(x-x_2)dt$ |
| Cell proliferation in State 2 | $x \mapsto x+1$ | $k_x x_2 dt$ |
| | $x_2 \mapsto x_2+1$ | |
| Cell switching from State 1 to 2 | $x_2 \mapsto x_2+1$ | $k_1(x-x_2)dt$ |
| Cell switching from State 2 to 1 | $x_2 \mapsto x_2-1$ | $k_2 x_2 dt$ |

Given that the event rates are linear functions of the state-space, the statistical moments of *x*(*t*), *x*_2_(*t*) can be obtained exactly using standard tools from moment dynamics. In particular, the time derivative of the expected value of any continuously differentiable function *ϕ*(*x, x*_2_) is given by

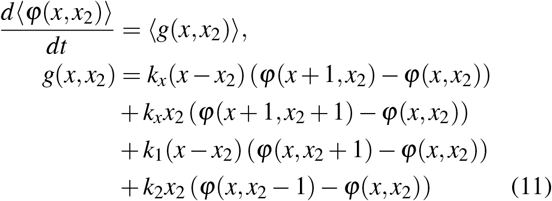

[59]–[62]. By setting *ϕ*(*x, x*_2_) = *x* and *x*_2_ in (11), one obtains the average population dynamics

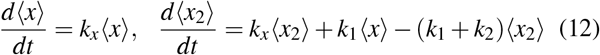

that is identical to the deterministically-formulated dynamics (3). Following the same steps for *ϕ*(*x, x*_2_) = *x*^2^, *xx*_2_ and 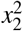

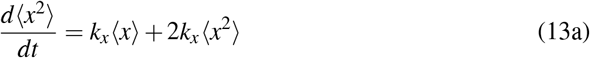

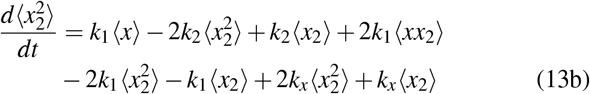

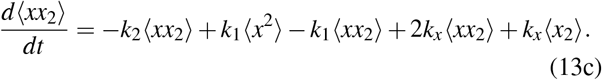

Based on the Bernoulli cell-state assignment of the starting cell, solving the linear dynamics system (12)-(13) with initial conditions

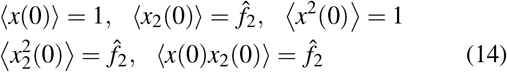

determines the statistical moments of *x*(*t*) and *x*_2_(*t*) at any time in the colony expansion. How do we use the moments of *x*(*t*) and *x*_2_(*t*) to obtain moments of their ratio *f*_2_(*t*) = *x*_2_(*t*)*/x*(*t*)? We next discuss two alternative approaches for doing this.

### C. Independent variable approximation

Analyzing the moment dynamic equations reveals

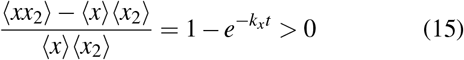

a positive covariance between the number of cells in State 2 and the colony size. Equation (15) points to weakly correlated *x* and *x*_2_ = *x f*_2_ for short times, implying an even weaker correlation between *x* and *f*_2_. This motivates an approximation where the fraction of cells in State 2 is independent of the colony size. Working along these lines, assuming *f*_2_(*t*) and *x*(*t*) to be independent allows an analytical derivation for the extent of fluctuations in *f*_2_(*t*) in spite of the nonlinear dependence. To see this, recall that by definition *x*_2_ = *f*_2_*x*, assuming ⟨*f*_2_*x*⟩ ≈ ⟨*f*_2_⟩ ⟨*x*⟩ and 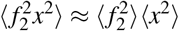, the moment of *f*_2_ can be obtained as

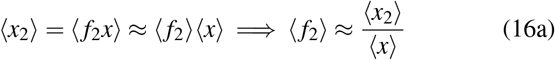

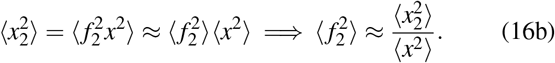

Substituting the solution of (12)-(13) in (16) results in 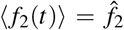 and the following formula for the extent of fluctuations

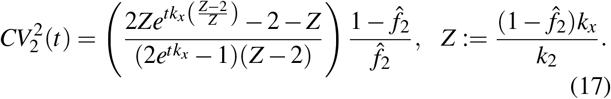

Note *Z >* 0 as 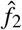 takes values between 0 and 1. When *Z* = 2, (17) reduces to

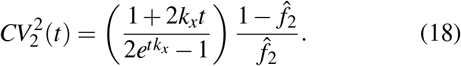

Here the dimensionless constants *tk*_*x*_ and *k*_*x*_*/k*_2_ have important biological interpretations: *tk*_*x*_ represents the average number of generations of colony expansion, and *k*_*x*_*/k*_2_ is the average number generations a cell remains in State 2 before switching to State 1. As expected from the Bernoulli initial condition,

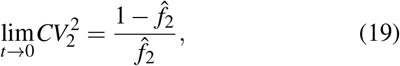

and if one grows the colony long enough, then 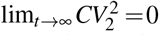 as *f*_2_(*t*) converges to 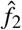in each colony. For a fixed 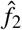 and *t, CV*_2_ monotonically increases with increasing time spent in State 2, i.e., slower switching results in higher fluctuations in the fraction of State 2 cells (Fig. 4).

**Fig. 4.**
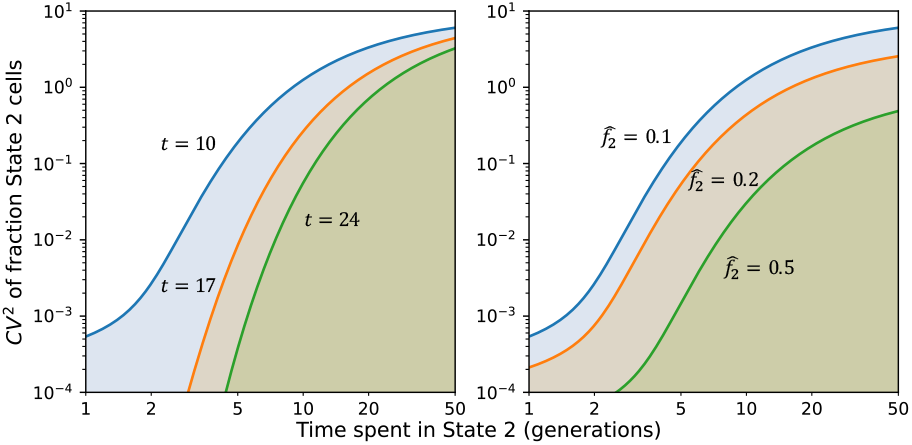
Inferring switching rates from the fluctuation test. *Left*: Colony-to-colony fluctuations in the fraction of State 2 cells as predicted by (17), as a function of time spent in State 2 for different durations *t* of colony expansion. In this plot, *k*_2_ is decreased to increase time spent in State 2, and *k*_1_ is adjusted to keep 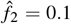. Thus, slower switching, where the memory of the initial state is retained and inherited for more generations, generates larger colony-to-colony fluctuations. *Right*: Same plot as on the left except *t* = 10 is fixed, and colony-to-colony fluctuations predicted by (17) are plotted for different fractions 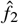. As before, *k*_*x*_ = 1, and *t* can be interpreted as time in the number of cell generations.

### D. Small noise approximation

An alternative derivation of *CV* relies on small fluctuations in *x*(*t*) and *x*_2_(*t*) around their respective means. Performing a Taylor series expansion of *f*_2_(*t*) yields

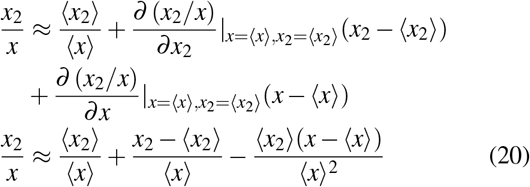

after ignoring quadratic and higher-order terms in the expansion. It is straightforward to see that in this limit

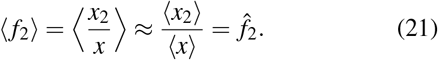

Moreover, from (20)

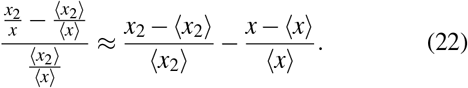

Squaring the terms on both sides and taking the expected value leads to

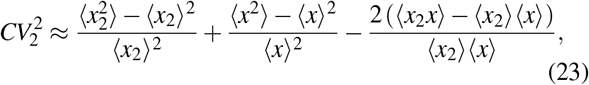

which after substituting the population number moments obtained from solving (12)-(13) yields the following formula

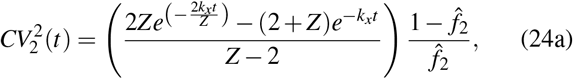

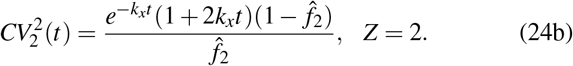

In summary, we have now developed three different formulas given by (10), (17) and (24), quantifying fluctuations in fraction State 2 cells across colonies. It is interesting to note that while all formulas reduce to (19) at *t* = 0, they have qualitatively different initial slopes

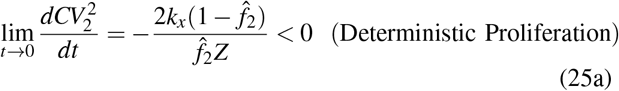

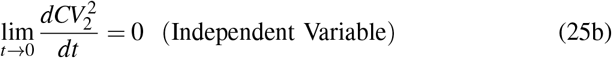

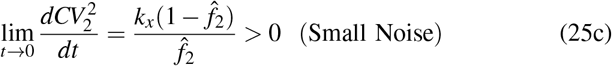

with fluctuations increasing with time in the early phase of colony expansion in (25c). All formulas share the common feature of lim_*t→*∞_ *CV*_2_ = 0.

Fig. 5 shows the accuracy of these formulas by comparison with exact *CV*_2_ values as obtained by performing stochastic simulations of the system shown in Table I. While the formula based on the deterministic formulation greatly underestimates fluctuations in *f*_2_(*t*), the independent variable assumption that takes into account stochastic proliferation provides a better approximation, especially at initial time points. The formula based on the small noise approach goes in the wrong direction at the start and overestimates the noise, but converges to the exact value at later time points when *CV*_2_ is small.

**Fig. 5.**
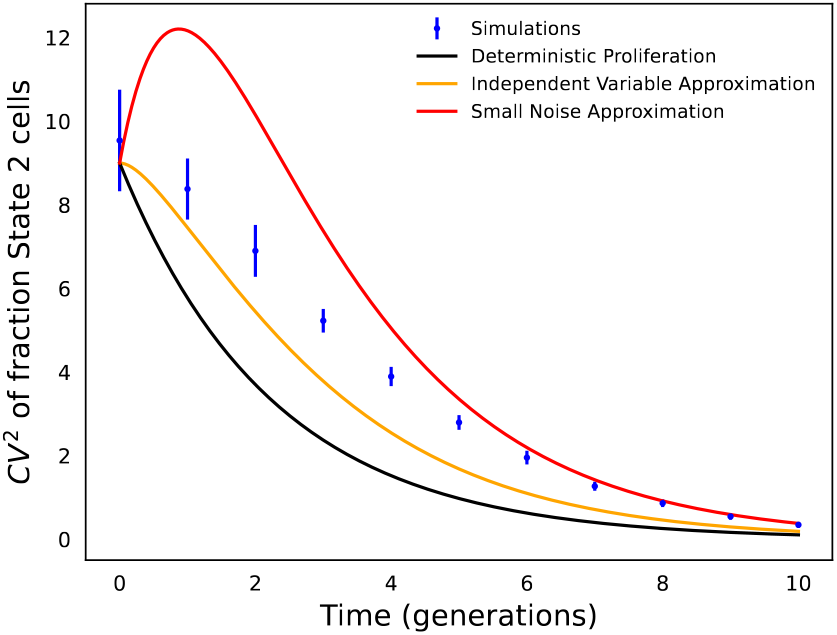
Comparison of analytical approaches to predict fluctuations in state fractions across single-cell colonies. Stochastic simulations of the model presented in Table 1 were used to simulate the fluctuation test experiment with 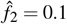, *k*_*x*_ = 1 and *k*_2_ = 1*/*5 (i.e., cells spend an average of 5 generations in State 2). *CV*_2_ is computed based on 1000 colonies, and simulations were repeated 20 times to generate error bars that show one standard deviation.

## IV. State transitions between multiple cell states

Having discussed an innovative approach to identify rates of switching between two states from measured inter-colony fluctuations, we now generalize this approach to multiple cell states. With more than two cell states there will be many different switching topologies, and one would like to identify plausible topologies from measured fluctuations in cell-state compositions. Here the fluctuation data will also be richer — one will measure both the variances and covariances in the fraction of different states across colonies. In this section, we show how to analytically predict these fluctuations for switching between an arbitrary number of cell states.

### Notation

Consider a system with *n* ≥ 2 cell states - States 1, 2, 3 …, *n*. In each state the cell proliferates with rate *k*_*x*_ and *k*_*i j*_, *i* ≠ *j* is the switching rate from state *i* to *j*. We denote by 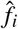 as the average fraction of cells in State *i* ∈ {1,…, *n*}. Random processes *x*_*i*_(*t*) and *f*_*i*_(*t*) represent the number and fraction of State *i* cells in the expanding colony, respectively, with total cell numbers *x*(*t*) = ∑ *x*_*i*_(*t*). Our goal is to quantify the colony-to-colony fluctuations in fraction State *i* cells

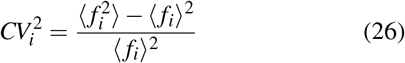

and the normalized covariances

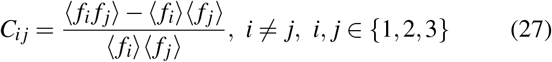

over time as a function of the proliferation/switching rates. Having described the notation used in this section, we briefly discuss the methodology for obtaining these fluctuations using the assumptions discussed in the previous section.

#### A. Deterministc proliferation Approximation

Let *f* (*t*) = [*f*_1_(*t*), …, *f*_*n*_(*t*)]^*T*^ denote a column vector of the fraction of cells in each state. Starting from a single cell that is in state *i* with probability 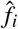, modeling colony expansion as an ordinary differential equation, *f* (*t*) evolves as per the linear dynamical system

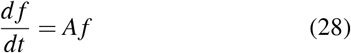

where elements of the *A* matrix are given by

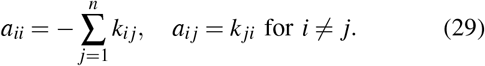

Solving (28) yields

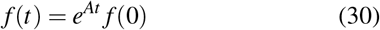

where the random vector *f* (0) is a column vector with zeros except 1 at the *i*^*th*^ row with probability 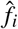, *i* ∈ {1,…, *n*}. Having obtained this initial-condition dependent random process *f* (*t*), (26) and (27) can directly be derived from it as done in (4)-(8).

#### B. Independent variable approximation

To account for stochastic proliferation one can build a model similar to Table I that enumerates all probabilistic events corresponding to cell proliferation in each state and switching between states. Let vector *μ* consists of all first- and second-order moments of integer-valued random processes *x*_1_(*t*), …, *x*_*n*_(*t*). Then, its time evolution can be derived as a linear dynamical system

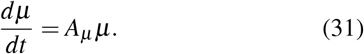

using moment dynamic tools [59], [61]. Solving this system with initial condition

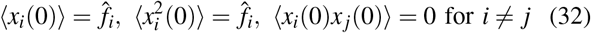

determines the time evolution of all the first- and second-order moments of cell numbers in different states. Having obtained these number moments, the moments of state fractions are obtained as

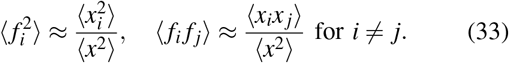

assuming that *f*_*i*_(*t*) is independent of *x*(*t*).

#### C. Small noise approximation

As derived in (23), for small deviations in cell numbers from their means the fluctuations in the fraction of State *i* is given by

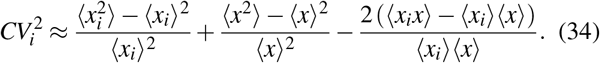

Using a similar Taylor series expansion approach it can also be shown that

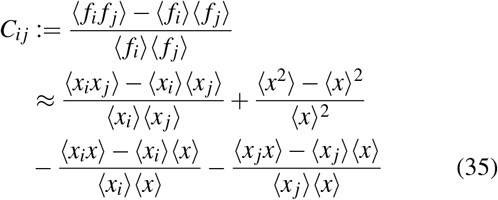

in the limit of small fluctuations in cell numbers. Substituting moments computed from (31) in (34) and (35) provides the extent of fluctuations and covariances between state fractions over time.

We illustrate these approximations on a ring topology of state switching where *k*_21_ = *k*_32_ = *k*_13_ = 0 (Fig. 6). Comparison with exactly-computed fluctuations from stochastic simulations shows that the independent variable approximation generally provides a good approximation at earlier time points when noise levels are high, while the small noise approximation works best at later time points.

**Fig. 6.**
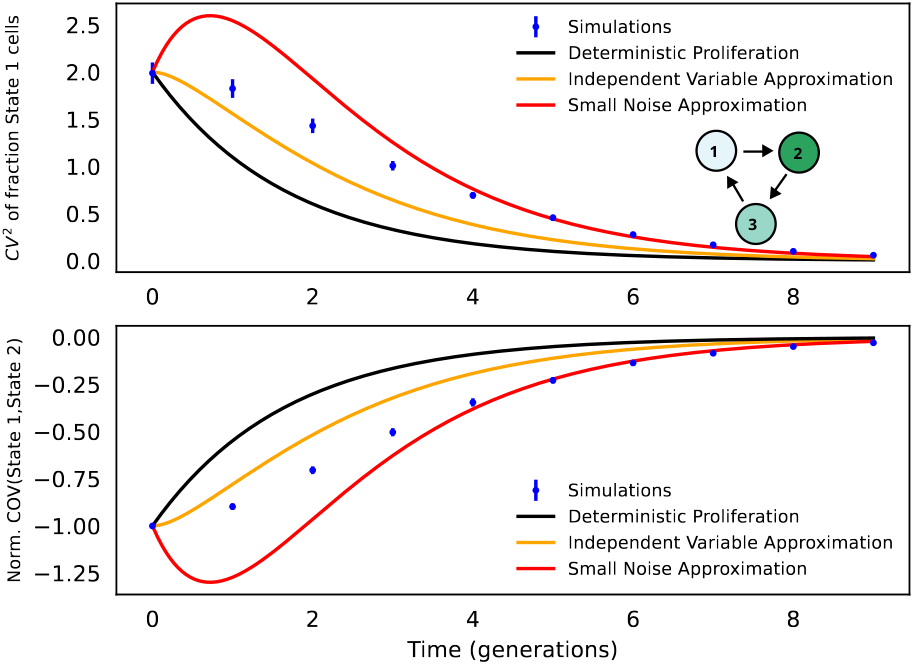
Extending the fluctuation test approach for switching between multiple states. Starting with a ring topology where *k*_21_ = *k*_32_ = *k*_13_ = 0 and *k*_12_ = *k*_23_ = *k*_31_ = 1*/*5, *k*_*x*_ = 1, 2000 colonies were simulated and plots show fluctuations in the fraction State 1 cells (top) and the normalized covariance between States 1 & 2 (bottom) over time. These fluctuations from stochastic simulations are compared with analytical formulas obtained using the three different approximations discussed in Section IV.

## V. Conclusion

The classical fluctuation test developed by Luria and Delbrück revolutionized the field of bacterial genetics and is employed to this day to estimate mutation rates (Fig. 1). While a genetic mutation is an *irreversible* transition, here we have used a similar fluctuation assay to probe reversible switching between cell states (Fig. 2). An advantage of this approach is that a single measurement of fluctuations in cell-state composition across colonies allows an estimation of a state’s transient heritability, i.e., how long a cell stays in a state before exiting it. While the approach only needs a single time point, performing the assay at several different time points is important for validating the inferred model. We have recently exploited his approach for diverse applications include characterizing drug-tolerant cancer cells [63]–[65], activation of human viruses such as HIV [66], and innate immune response in individual human cells [67].

The key contribution of this work has been the development of different analytical formulas for quantifying the extent of inter-colony fluctuations. While the formula based on deterministic proliferation is the simplest, it may result in inferred switching rates that are slower than their actual values as it underestimates fluctuations (Fig. 3). Using formulas based on stochastic proliferation may provide more accurate inference. Future work will consider expanding these formulas to scenarios where states have different proliferative potentials and this is especially important given that drug persisters can in some cases grow significantly slower than drug-sensitive cells [68]–[73].

We further generalized these analytical formulations to switching between an arbitrary number of cells states. While here we have primarily focused on the forward prediction of fluctuations given a switching topology, the backward inference problem of using measured fluctuation signatures to find the most parsimonious switching topology provides fertile grounds for future work.

